# Double Dissociation of Nicotinic α7 and α4/β2 Sub-receptor Agonists for Enhancing Learning and Attentional Filtering of Distraction

**DOI:** 10.1101/369496

**Authors:** Maryzeh Azimi, Mariann Oemisch, Thilo Womelsdorf

## Abstract

Nicotinic acetylcholine receptors (nAChR) modulate attention, memory, and higher executive functioning, but it has remained unclear whether nAChR sub-receptors tap into different neural mechanisms of these functions. We therefore set out to contrast the contributions of selective alpha-7 nAChR and alpha-4/beta-2 nAChR agonists in mediating value learning and attentional filtering of distractors in the nonhuman primate. We found that the alpha-7 nAChR agonist PHA-543613 selectively enhanced the learning speed of feature values but did not modulate how salient distracting information was filtered from ongoing choice processes. In contrast, the selective alpha-4/beta-2 nAChR agonist ABT-089 did not affect learning speed but reduced distractibility. This double dissociation was dose-dependent and evident in the absence of systematic changes in overall performance, reward intake, motivation to perform the task, perseveration tendencies, or reaction times. These results suggest nicotinic sub-receptor-specific mechanisms consistent with (1) alpha-4/beta-2 nAChR specific amplification of cholinergic transients in prefrontal cortex linked to enhanced cue detection in light of interferences, and (2) alpha-7 nAChR specific activation prolonging cholinergic transients, which could facilitate subjects to follow-through with newly established attentional strategies when outcome contingencies change. These insights will be critical for developing function-specific drugs alleviating attention and learning deficits in neuro-psychiatric diseases.

## Introduction

Nicotinic acetylcholinergic receptors (nAChR’s) have long been implicated to modulate attention and learning (Deutsch JA 1971; Muir JL et al. 1992), and are primary targets for developing drug treatments for reinstating attention and flexibility to adjust behavior in schizophrenia, attention deficit hyperactivity disorder (ADHD) and Alzheimer’s disease (Wilens TE et al. 1999; Wallace TL, TM Ballard, et al. 2011; Ballinger EC et al. 2016). These effects are mediated by the prefrontal cortex whose innervation from basal forebrain cholinergic neurons is essential for successful reversal learning (Roberts AC et al. 1990; Wallace TL and D Bertrand 2013), and whose cholinergic concentration levels predict successful attentional detection of relevant visual cues (St Peters M et al. 2011; Howe WM et al. 2013). These insights into the cholinergic contribution to successful attention and learning raises the long-standing question whether these functional contributions can be dissociated, or else, might reflect the same underlying processes (Sarter M et al. 2003).

Evidence from a variety of sources suggests that the cholinergic influence on attention and learning functions might be mediated differentially by the α7 and the α4β2 nAChR sub-receptor systems (Sarter M et al. 2016; Thiele A and MA Bellgrove 2018). Both sub-receptors are expressed in prefrontal cortex and act pre-synaptically as autoreceptors or heteroreceptors to regulate the release of glutamate, GABA, dopamine and other neurotransmitters (Gotti C et al. 2006; Livingstone PD and S Wonnacott 2009; Bortz DM et al. 2013). However, subtle differences exist in their permeability of calcium and associated role in plasticity (stronger for α7), and in the amplification potential of glutamate signaling (stronger but shorter lasting for α4β2) (Albuquerque EX et al. 2009; Parikh V et al. 2010; Tanner JA et al. 2015). Moreover, prefrontal α7 and α4β2 receptors show a layer-specific expression profile with stronger α4β2 expression in thalamic recipient layer VI and α7 more prominent expression in layer V, which is rich in striatal projection neurons (Poorthuis RB et al. 2013). These differences in layer-specific action, calcium-mediated plasticity, and control of glutamate excitability make it possible that each sub-receptor plays separable circuit roles for attention and learning functions of the prefrontal cortex. To discern dissociable circuit roles, it is essential to establish a task paradigm and animal model showing sub-receptor specific sensitivity to attention and learning functions.

Task paradigms that simultaneously vary attentional demands and are sensitive to learning processes are sparse, which constitutes a bottleneck for advancing a mechanistic understanding of sub-receptor specific functions (Sarter M *et al.* 2003; Romberg C et al. 2013). For instance, in NHP studies, investigation of attentional processes in the context of nicotinergic experiments have been largely limited to testing distracting stimuli in working memory tasks (surveyed in **Table S1, S2**). To address this, we selected two sub-receptor agonists and tested them on a feature-based attention reversal learning paradigm in nonhuman primates (NHP’s) (**Figure 1**). The task distinguished feature-based covert attention from spatial response biases, varied the demands of attentional filtering of distractors, and quantified reversal learning flexibility. We conjectured that such a combined learning and attention paradigm is needed to clarify open questions about sub-receptor specific functions. In particular, an exhaustive literature survey of attention effects of fourteen studies applying systemically α7 agonists (**Table S1**) (Briggs CA et al. 1997; Hahn B et al. 2003; Buccafusco JJ et al. 2007; Buccafusco JJ and AV Terry, Jr. 2009; Rezvani AH et al. 2009; Wallace TL, PM Callahan, et al. 2011; McLean SL et al. 2012; Gould RW et al. 2013; Yang Y et al. 2013; Young JW et al. 2013; Jones KM et al. 2014; Kolisnyk B et al. 2015; Wood C et al. 2016; Wadenberg MG et al. 2017), and thirteen studies using α4β2 agonists (**Table S2**) (Buccafusco JJ et al. 1995; Decker MW et al. 1997; Prendergast MA, AV Terry, Jr., et al. 1998; Schneider JS et al. 1999; Schneider JS et al. 2003; Decamp E and JS Schneider 2006; Buccafusco JJ *et al.* 2007; Howe WM et al. 2010; Gould RW *et al.* 2013; Paolone G et al. 2013; Kolisnyk B *et al.* 2015; Terry AV, Jr. et al. 2016; Wood C *et al.* 2016) failed to reveal sub-receptor specific contributions to working memory and distractor filtering functions in the NHP animal model.

**Figure 1.**
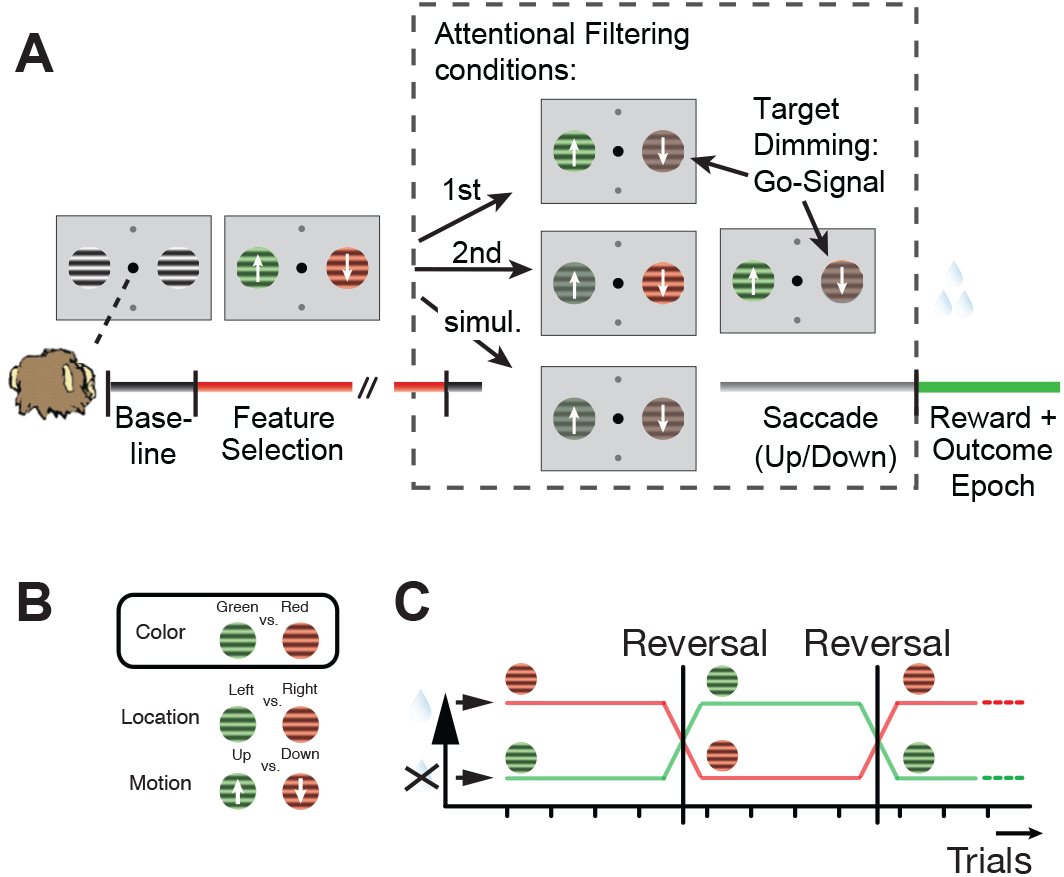
Feature-based reversal learning task. (**A**) The task requires centrally fixating and covertly attending one of two grating stimuli that has the color associated with reward. To obtain reward the animal had to saccade in the up-/downward direction of the motion direction of the attended stimulus. The saccadic choice had to be made within 0.5 s following a dimming of the target stimulus. The dimming served as a Go-signal for the saccadic choice. In three attentional filtering conditions, the target Go-signal could either occur (1) in the target first, i.e. before the distractor, (2) in the target stimulus second, after the dimming occurred in the distractor stimulus, or (3) the dimming event occurred simultaneously in target and distractor (indicated in dashed box). (**B**) Three features characterize each stimulus – color, location, and motion direction. Only the color feature is directly and consistently linked to reward outcome. (**C**) The task is a reversal learning task, whereby only one color at a time is rewarded. This reward contingency switches repeatedly and unannounced across blocks of trials.

A similarly inconclusive pattern of results exists for set shifting and reversal learning tasks. Studies in rodents indicate that attentional set shifting and behavioral flexibility is closely associated with α7-specific signaling across five independent studies amplifying α7 sub-receptor activation with different systemic agonists (McLean SL et al. 2011; Wallace TL, PM Callahan*, et al.* 2011; McLean SL *et al.* 2012; Jones KM *et al.* 2014; Wood C *et al.* 2016; Wadenberg MG *et al.* 2017) (**Table 1** and **Table S1**). However, in the one rodent study that compared a β agonist to an α7 agonist, similar set-shifting behavioral benefits were found with both sub-receptor specific types (Wood C *et al.* 2016). This result resonates with findings in the NHP where a study with mPTP-impaired NHPs reported that α4β2 agonist action was sufficient to normalize spatial reversal learning performance (Decamp E and JS Schneider 2006) in that both studies showed beneficial effects of selective agonists on a task component involving reversal of feature values. However, in conflict to these studies one study reported slower object reversal learning in NHP’s with a selective α7 agonist applied at a dose at which it improved delayed match-to-sample performance in the same monkeys (Gould RW *et al.* 2013), suggesting that α7 agonists can have negative influences on reversal learning at doses that working memory performance would benefit from.

**Table 1.**
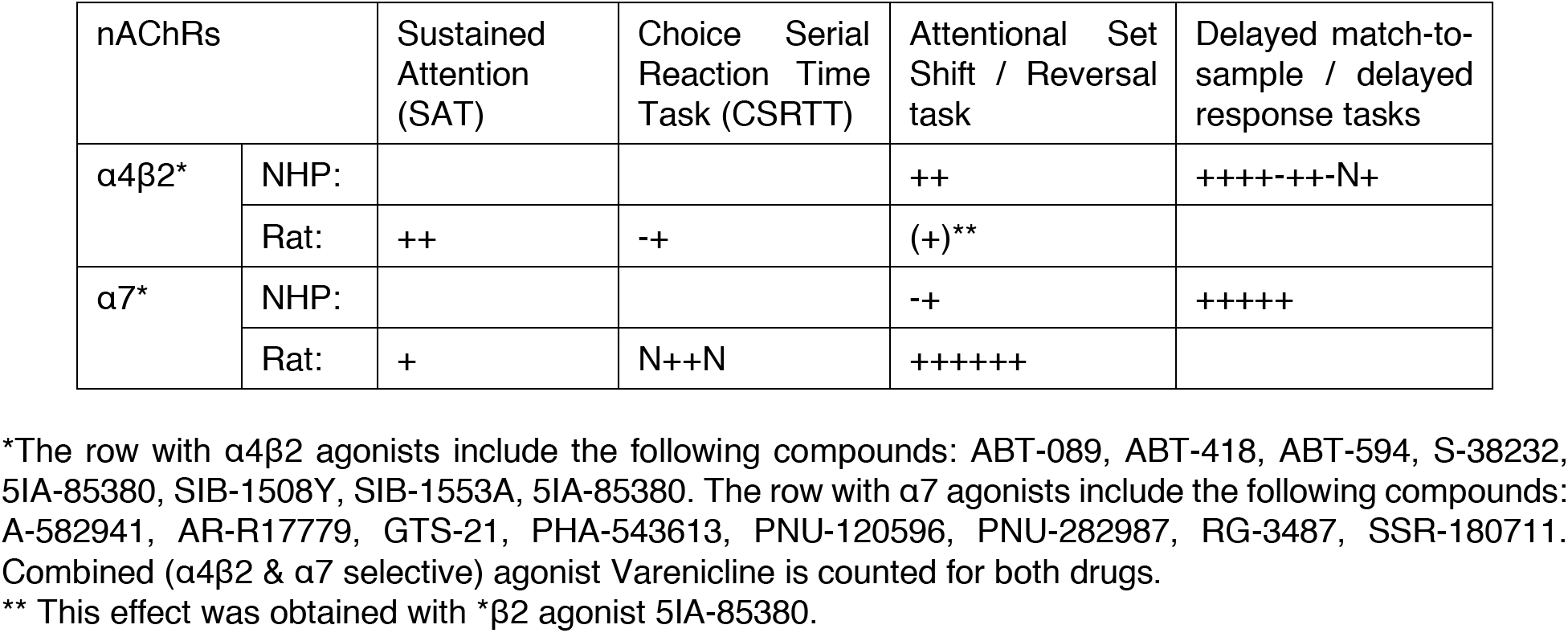
Overview of results from studies testing the influence of systemic application of selective α4β2 and α7 nAChR’s on attention related performance in rodent and nonhuman primate (NHP’s). The details about the fourteen studies using α7* and thirteen α4β2* studies included are outlined in **Table S1** and **Table S2**. The tasks include the sustained attention task (SAT), the choice serial reaction time task (CSRTT), in attentional set shifting and reversal tasks, and in delayed response tasks including the delayed-match-sample task. The symbols denote the result of individual studies reporting either (1) enhanced for at least one dose (“+”), or (2) or only reduced performance (“-”), or which report no effect (“N”).

In the context of the described result patterns, we aimed to establish for the NHP an attention and learning paradigm and identifies independently for an alpha7 and an α4β2 agonist the dose with likely beneficial behavioral effects. We report that α7 agonists were associated with faster reversal learning, that α4β2 agonists were associated with enhanced performance when distractor filtering was most demanding, and that neither sub-receptor system affected motor, reward, or overall performance components of the task.

## Materials and Methods

Data was collected from a 9 and a 7-year-old adult male rhesus monkey (*Macaca mulatta*) following procedures described in (Hassani SA et al. 2017). All animal care and experimental protocols were approved by the York University Council on Animal Care and were in accordance with the Canadian Council on Animal Care guidelines.

### Behavioral Procedures and Paradigm

During experiments monkeys were seated inside a primate chair, head-fixed and 58 cm away from a 21-inch LCD monitor inside a sound attenuating chamber. Visual stimuli, gaze (SRS Eye-link 1000 system), and fluid reward delivery were controlled by MonkeyLogic toolbox (http://www.brown.edu/Research/monkeylogic/). The monkeys performed a feature-based reversal learning task that required covert spatial attention to one of two stimuli dependent on color-reward associations (**Fig 1A**). In contrast to color, other stimulus features (motion direction and stimulus location) were only randomly related to reward outcome – they were pseudo-randomly assigned on every trial (**Fig 1B**). Color-reward associations were reversed in an uncued manner between blocks of trials with constant color-reward association (**Fig 1C**). Each trial started with the appearance of a grey central point, which had to be fixated. After 0.5 - 0.9s, two 2° radius wide black/white gratings appeared 5° to the left and right of the central fixation point with 0.8 °/s motion inside the apertures drifting in opposite directions. Following another 0.4s the two stimulus gratings either changed color to black/green and black/red (Monkey K: black/cyan and black/yellow), or started moving at 1.20 cycles/degree in opposite directions up and down, followed after 0.5 - 0.9s by the onset of the second stimulus feature that had not been presented so far, e.g. if after 0.4s the grating stimuli changed color then after another 0.5 - 0.9s they started moving in opposite directions. Within the next 0.4 - 1s either the red and green stimulus dimmed for 0.3s followed by the dimming of the other stimulus after 0.55 s, or they dimmed simultaneously. The dimming represented the Go-cue to make a saccade to one of two response targets displayed 4° above and below the central fixation point. The monkeys kept central fixation until this dimming event occurred. A saccadic response following the dimming was only rewarded (with ~0.3ml fluid) if it was made in the direction of motion of the stimulus carrying the reward associated color and at the time that stimulus dimmed. The color-reward association was changed every 30-50 trials and 30-100 trials for Monkey H and Monkey K respectively. During task performance, a running average of 90% rewarded trials over the last 12 trials induced an un-cued block change.

### Experimental Procedure

Monkey H and Monkey K did not receive either of the two drugs prior to the experiment. For each monkey, prior to the first week with drug/saline injections we obtained baseline performance. Monkey H started with PHA-543613, while monkey K started with the ABT-089 drug conditions before the drugs were reversed. Data was collected from Tuesday to Friday with two days pseudo-randomly assigned to drug injection (IM) and the remaining two days assigned to vehicle (saline) injections (IM). The assignments of drug days balanced the number of days of the week between drug and control condition to preclude prediction of when drugs versus vehicles were injected. The vehicle injections occurring between drug treatments were considered for analyses as control sessions. Drugs and vehicle were injected by a lab technician. The experimenter was blind to the treatment conditions. For each experimental session, subjects were given at least 50 minutes to perform the task. For PHA-543613 experiment, injection was administered 30±1 minute prior to start of the task. The time frame was chosen as an estimate based on prior (mostly rodent) studies (Wishka DG et al. 2006; Yang Y *et al.* 2013; Bali ZK et al. 2015; Kolisnyk B *et al.* 2015; Sadigh-Eteghad S et al. 2015). For ABT-089 experiment, the injection was done 10±1 minutes prior to the start of the task. This time frame was chosen based on previous non-human primate studies where the same drug was applied (Decker MW *et al.* 1997; Prendergast MA, WJ Jackson, et al. 1998). All sessions were conducted at the same time of the day. The selected doses for PHA-543613 experiment with Monkey H were 0.125 and 0.250 mg/kg. Once the optimal dose was determined for Monkey H, only the 0.250 mg/kg dose was used with Monkey K. These doses were chosen as an estimate from the rodent literature (Wishka DG *et al.* 2006; Bali ZK *et al.* 2015; Kolisnyk B *et al.* 2015; Sadigh-Eteghad S *et al.* 2015). The doses for the ABT-089 experiment (0.01and 0.02 mg/kg) were chosen based on high and low values of the dose range previously tested in rhesus monkeys (ref). Initially, 0.04 mg/kg was selected as the higher dose for Monkey H, but discontinued after possible adverse side effects (nausea) could not be excluded. Once, the optimal dose was determined for Monkey H, the 0.02 mg/kg dose was used with Monkey K, however, upon observation of no improvement in the behavior and a slight tendency towards worse performance in one of the task components, the 0.01 mg/kg dose was administered to Monkey K. Experiments with PHA-543613 and ABT-089 could be analyzed for 8 and 12-week for monkey H, and for 2 and 4 weeks for monkey K. For Monkey H, we analyzed 15, 8, and 7 sessions for control, 0.250 mg/kg and 0.125 mg/kg dose of PHA-543613, respectively, and 20, 7, and 9 sessions for control, 0.02 mg/kg and 0.01 mg/kg dose of ABT-089, respectively. For Monkey K, we analyzed 7, 4, and 4 sessions for control, 0.02 mg/kg and 0.01 mg/kg dose of ABT-089, respectively, and 3 and 4 sessions for control and 0.25 mg.kg for PHA-543613, respectively.

### High Pressure Liquid Chromatography (HPLC) Analysis of Blood Serum

We used mass spectrometry HPLC analysis to identify the peak concentration and overall metabolism pattern of the drugs over time. Nine blood samples of 300 μl in total (baseline and 8 samples after injection) were extracted from a 10-year-old, 10 kg male rhesus macaque. The samples were taken at the following time points: 1 minute before drug injection (baseline), 9,15,25,40,70,100,160 & 220 minutes after injection. The higher drug dose (0.25 mg/kg) for PHA-534613 was considered for HPLC analysis while for ABT-089 the lower dose (0.01 mg/kg) was selected. Samples were centrifuged at 2000 rpm speed for about 40 minutes and after being spin filtered, stored for mass spectrometry HPLC processing. The procedure was successful for PHA-543613 (**Fig. S1**)., However, no AB-089 signal was observed in any of the blood samples and as a result no blood serum analysis could be obtained. It was possible that the injected drug dose was too low or the drug became completely protein bound in the blood and therefore would not have been able to pass through the spin filter. In previous literature, the plasma exposure of ABT-089 was measured in baboons under both bolus intravenous (IV) injection and slow infusion in doses ranging 0.04-1 mg/kg (Chin CL et al. 2011).

On average, Monkey H worked for 74 and 60 minutes and Monkey K worked for 84 and 92 minutes during experimental sessions of PHA-543613 and ABT-089, respectively. Results from (Chin CL *et al.* 2011) and HPLC analysis of PHA-543613 suggested that the peak concentration and the subsequent drop in blood concentration were captured within the testing session time for both pharmacological agents.

### Data Analysis

Analysis was done using Matlab (The Mathworks). To assess the effects of systematic injection of PHA-543613 and ABT-089 on attention and feature based reversal learning, different aspects of behavior were defined and quantified as described below. In some analyses, a moderately small number of data points existed in the drug condition which was considerably fewer than control. Therefore, data across all control and drug sessions were compared using non-parametric Wilcoxon rank sum test for evaluation of performance in different time windows, trial type subsets based on dimming event and number of reversal blocks and rewarded trials. In addition, two-sample z-test was used for comparing proportions and randomization test was applied when correction for multiple comparison was needed.

### Temporal variation of drug effects on performance

Blocks were categorized as learned and not-learned; the latter defined as blocks with no identified learning trials (the first trial which the ideal observer component of the model identifies for reliable learning to occur above chance performance) or having learning trials above 24 as computed by the expected maximization (EM). Performance for the analysis of temporal variation of drug effects was only done in learned reversal blocks. Trial by trial performance was estimated by using an Expectation Maximization algorithm (Smith AC et al. 2004). For details on implementation of EM algorithm in modelling the performance of subjects, see (Hassani SA *et al.* 2017). We tested drug effects at different time windows relative to the injection of the drug similar to previous studies of selective nAChR agonists with different plasma level half times to prevent overlooking shorter lasting effects (e.g. (Hahn B *et al.* 2003). Different overlapping time windows after the task started were selected to evaluate the temporal effects of drugs on behavior. The time windows included: 0-25, 12.5-37.5, 25-50, 37.5-62.5 minutes. **
Table 3** and **Table 4** show the number of learned blocks within each time window and during the entire session across all treatment conditions for Monkey H and Monkey K, respectively.

Dynamics of performance including learning speed and net increases in performance were measured via parameters of a hyperbolic ratio function fit to learning curves. The parameters of interest to compare between experimental and control sessions were C_50_ (trial to reach half maximal of performance, exponent (slope) and R_max_ (maximum increase in performance from baseline performance). Randomization procedure was carried out to test the significance of differences in the parameters of interest estimated by the hyperbolic-ratio function between drug and control conditions. The randomization procedure followed steps described in (Maris E and R Oostenveld 2007). The randomization was repeated 1000 times and the distribution of root of mean square error (RSME) of all estimated parameters was constructed. Any estimation that had RSME above 90 percentile of this distribution was excluded. The difference between the estimated parameter values were extracted as test statistics and the 97.5% and 2.5% percentiles of the null test statistics distribution was computed to conduct a two-tailed test at 0.05 alpha significance. These two values served as thresholds based on which the significance of the observed difference between the parameters of interest was determined. The proportion of values within the null distribution larger than the observed test statistics was calculated as the p value of this randomization procedure.

### Distractor filtering

Distracting effects of irrelevant stimuli were evaluated in trials were dimming of both stimuli occurred simultaneously. Here, performance was calculated as the proportion of correct choices across all blocks of a session. In comparison to trials where dimming of the two stimuli did not happen simultaneously, performance in the same dimming trial type required a higher level of attentiveness to filter distraction. All dimming trial types were presented with the same ratio i.e. 1/3 each.

### Motivation

The motivation of subjects was studied in terms of total number of blocks performed and rewarded trials which would indicate the amount of earned fluid during the sessions. The proportion of learned to total reversal blocks were computed as well and compared to each other with two sample proportions z-test. The z score was computed based on the equation below (Zar JH 2010) where *p*_1_ is the first proportion value, *p*_2_ is the second proportion value. If the null hypothesis was true, then *p*_1_-*p*_2_=0. *P* is the proportion of learned blocks calculated by pooling data from both control and drug conditions as shown by the following equation:

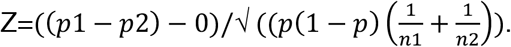

## Results

We obtained feature-based learning performance with a α7-nAChR agonist PHA-543613 in 19 sessions (monkey H= 8 sessions for 0.250 mg/kg and 7 sessions for 0.125 mg/kg dose), monkey K=4 sessions for 0.250 mg/kg; saline control sessions: 15/3 for monkey H and K, respectively), and with a α4β2-nAChR agonist ABT-089 in 24 sessions (monkey H= 7 sessions for 0.02 mg/kg and 9 sessions for 0.01 mg/kg dose, monkey K=4 sessions for 0.02 mg/kg and 4 sessions for 0.01 mg/kg; saline control sessions: 20/ 7 for H and K, respectively).

## Overall Performance Effects of Selective nAChR

Both nAChR agonists had a negligible influence on overall performance accuracy and motivation to perform the task. For PHA-543613 the overall performance accuracy was 77%, 77%, and 74% percent correct for the 0.25 mg/kg, 0.125 mg/kg and control conditions in monkey H, and 65% and 63% percent correct for the 0.25 mg/kg and control conditions in monkey K. The different performance levels were not significantly different (for monkey H and K, all tests, z test, p>0.05). For ABT-089 the overall performance accuracy was 83%, 83%, and 80% percent correct for the 0.02 mg/kg, 0.01 mg/kg, and control conditions in monkey H, and 63%, 69%, and 65% percent correct for the 0.02 mg/kg, 0.01 mg/kg and control conditions in monkey K. The different performance levels were not significantly different (for monkey H and K, all tests, z test, p>0.05).

## nAChR Agonists Effects on Motivation and Reward Intake

We estimated the motivation to perform the task by calculating the number of reversal blocks the animals engaged in across sessions with and without drug. For PHA-543613, we found that the number of reversal blocks performed during 0.25 mg/kg, 0.125 mg/kg, and control sessions were not significantly different for monkey H, or monkey K (all Wilcoxon ranksum tests, p>0.05). Similarly, for ABT-089, we found that the number of reversal blocks performed during 0.01 mg/kg, and control sessions were not significantly different for monkey H, or monkey K (all Wilcoxon ranksum tests, p>0.05). The only exception was for the high (0.02 mg/kg) dose for which monkey H performed fewer number of blocks compared to control sessions (Wilcoxon ranksum test, p=0.0255). However, during these sessions for monkey H the proportion of learned versus unlearned blocks with ABT-089 (0.02 mg/kg) treatment was significantly higher than control (z test, p=0.018). This higher proportion of learned to unlearned blocks could account for the lower number of reversal blocks performed in the same sessions and are unlikely a reflection of drug effects on motivation.

Consistent with the previous results, we found that the overall reward intake per session, calculated as the total number of rewarded choices per session, was similar between drug and control experiments for both drugs. For PHA-543613 the average number of rewarded trials per session in the 0.250 mg/kg, 0.125 mg/kg and control conditions was 342+/-30.47^1^, 401+/-61.99 and 354+/-26.51 for monkey H, and 245+/-14.04 and 248+/-13.42 for monkey K (all Wilcoxon ranksum tests, p>0.05). For ABT-089 the average number of rewarded trials per session in the 0.02 mg/kg, 0.01 mg/kg and control conditions was 212+/-17.13, 304+/-35.32 and 264+/-17.79 for monkey H and 216+/-21.86, 189+/-26.36 and 205+/-21.90 for monkey K (all Wilcoxon ranksum tests, p>0.05). Taken together, neither nAChR drug modulated overall motivation and reward intake at the doses tested.

## Effects of nAChR Agonists on Feature-based Learning

We next calculated whether the drug treatment facilitated learning of feature values. To this end, we computed the learning curves for each reversal block using an ideal observer statistics that calculated how consistently the monkeys choices were rewarded across sequences of trials (*see* Methods, and (Hassani SA *et al.* 2017)). We then fit the resulting learning curves with a hyperbolic ratio function to extract the trial number at which the learning reached 50% of the maximum asymptotic performance value, which could be early or late, corresponding to fast versus slow learning (**Fig. S2**). We term this learning trial the “L_50_“ (in analogy to the C_50_ for contrast sensitivity measurements using the same function). L_50_’s for the overall learning curves did not differ for either monkey, dose and drug condition, as expected from the similar overall performance (*see* above). Given the known temporal decay of the receptor agonists over the course of 1-2 h sessions (see Methods and **Fig. S1**), we thus compared the distribution of L_50_’s for learning curves obtained in successive 25 minute time intervals following systemic drug or saline injections.

For the PHA-543613 treatment, we found that learning speed as indexed by the L_50_ parameters were not different between the 0.125 mg/kg dose and control injections for Monkey H in either 25 minute time window after drug injection (randomization test, p>0.5) (**Fig. 2A,B**). However, both monkeys showed faster learning speeds with the higher dose during specific time periods with 0.25 mg/kg PHA-543613 compared to the saline control injections. At this higher dose, monkey H showed in the first 25 minute a significantly earlier average L_50_ in the drug versus control conditions (randomization test, p=0.0341) (**Fig. 2A, left panel**). No later time epoch showed different average L_50_’s (randomization tests all n.s.). Monkey K showed a significant earlier average L_50_ after drug versus control injection conditions in T2 (12.5-37.5 minute interval) (randomization, p=0.022) and in the T3 (25-50 minute time interval) (randomization, p=0.0123) (**Fig. 2B, left panel**). No other time epoch showed an L_50_ effect for drug versus control conditions (randomization tests all n.s.).

**Figure 2.**
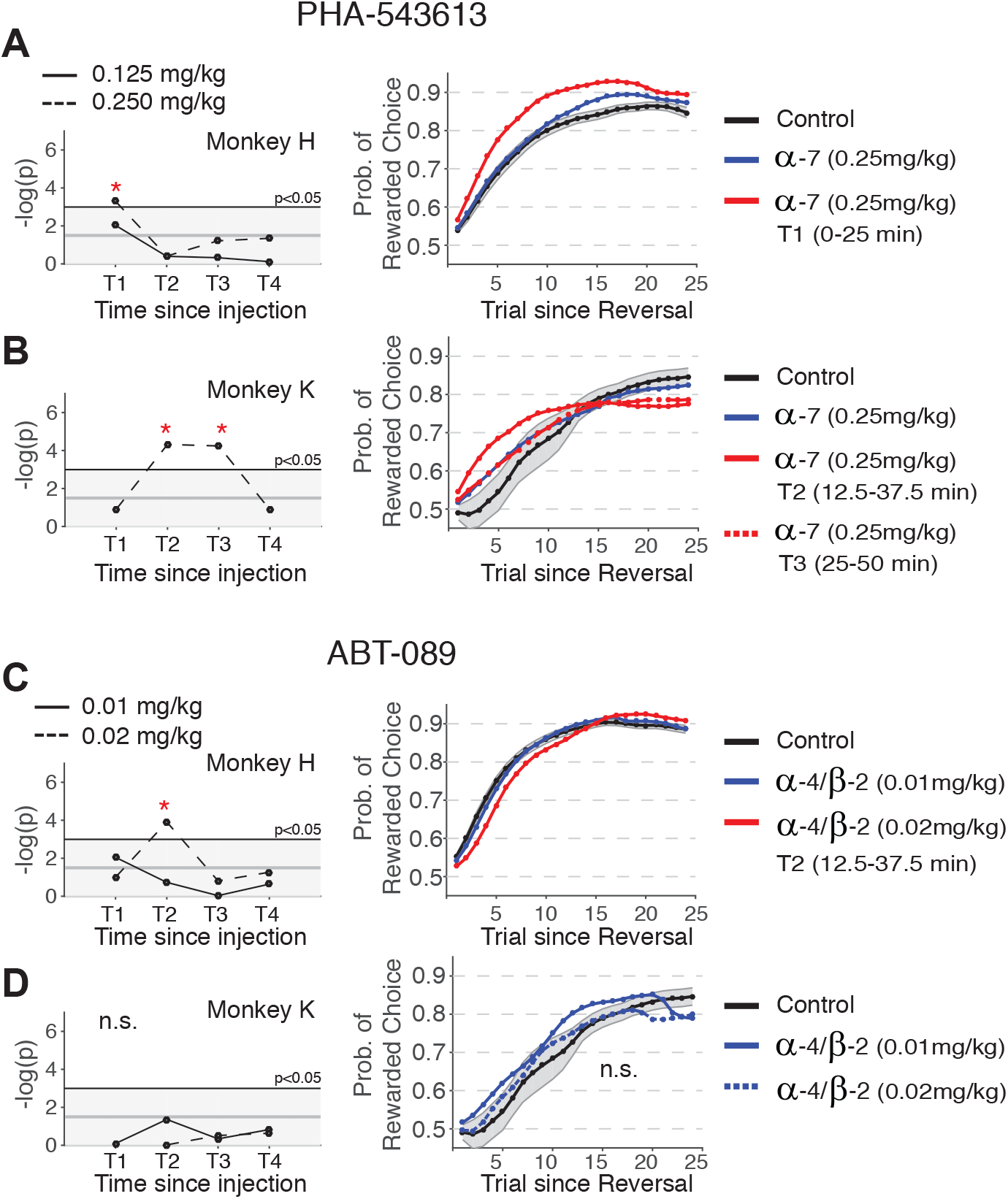
Sub-receptor specific effects on reversal learning speed. (**A**) *Left:* Distribution of p values from Bootstrap analysis of learning speed at four different time windows from the time of injection of PHA-543613 or vehicle for Monkey H. The solid (dashed) line denotes the statistical comparison of learning speed between 0.125 (0.250) mg/kg PHA-543613 and the vehicle control condition. P values larger than three (-log(0.05)) denote a significantly different learning speed between drug and control condition. Time windows T1 to T4 refer to 0-25, 12.5-37.5, 25-50 and 37.5-62.5-minute time windows relative to the beginning of the testing session. Testing sessions started 30 min. and 10 min. following drug/vehicle injection during PHA-543613 and ABT-089 experiments, respectively. *Right:* The learning curves for Monkey H across control performance (black), the average performance in the 0.25mg/kg PHA condition (blue), and within the time window T1 (red), which revealed a significant faster learning in the bootstrap analysis (see left panel). (**B**) Same format as *A* for Monkey K, showing significant faster learning with 0.250 mg/kg PHA-543613 in the time windows T2 and T3 (red solid and dashed lines). (**C**) Same format as *A* for Monkey H and injection of ABT-089, showing no learning improvement, but a significantly decreased learning speed at time window T2 with the lower (0.01 mg/kg) dose of ABT-089. (**D**) Same format as *C* for Monkey K, showing no significant effect of ABT-089 on learning speed at any time window. Monkey K’s baseline vehicle control condition is from the PHA-543613 control (?) sessions. Error bars denote standard error of the mean.

In contrast to the PHA-543613 treatment effects, ABT-089 did not increase the learning speed at any 25 minute time interval following injection (**Fig. 2C,D**). Across all statistical comparisons, we found only for the high 0.02mg/kg ABT-089 dose a significant modulation at the T2 (12.5-37.5 minute time interval), evident in slower learning in the drug condition (mean L_50_ of 6.02 vs. 4.34 for drug and control condition, respectively; Bootstrap significance p=0.02 **Fig. 2C**, *right panel*). At all other conditions the L_50_’s in drug and control conditions were indistinguishable (for monkey’s H and K, all comparisons n.s.; the 0.02 mg/kg dose, T1 (0-25 minute time interval) in monkey K was not tested because too few learned blocks were available for analysis).

In addition to the L_50_, we also characterized the plateau level of performance of the learning curves using the R_max_ parameter of the hyperbolic ratio fits and found inconsistent results. Across the treatment with PHA-543613 or ABT-089, at low and high doses, and at either of four different time epochs since injection we found that for Monkey H, R_max_ was lower at T1 (0-25 minute window) after 0.02 mg/kg ABT-089 injection (randomization test, p=0.0311). For Monkey K, R_max_ was lower in PHA-543613 0.250 mg/kg condition than control condition during the T2 (12.5-37.5 minute window) (randomization test, p=0.0379 Fig. S5-A, *left panel*). In the same time window, ABT-089 0.01 mg/kg led to higher R_max_ performance (randomization test, p = 0.0174).

## Effects of nAChR Agonists on Attentional Filtering of distraction

Our feature-based reversal learning task required monkeys to make a saccadic choice (up-/downwards saccade) to a transient dimming event of the stimulus with the reward associated color. We used this choice event to manipulate the attentional filtering demands of the task by having both stimuli dimming at the same time in one-third of the trials (**Fig. 1**). In these trials, the dimming of the distractor stimulus had to be attentionally filtered in order to process the dimming of the target stimulus that had the reward associated color. In the other two thirds of trials the target stimulus dimmed before or after the dimming of the distractor stimulus, and thus required filtering the dimming event in the distractor without concomitant processing of a dimming event of the target. To compare the drug effects on attentional filtering we compared the proportion of erroneous choices to the motion direction of the distractor separately for the condition with the target dimming before, after, and simultaneously with the distractor (**Fig. 3A,C**; for detailed data for each monkey and subcondition, please see **Fig. S3**).

**Figure 3.**
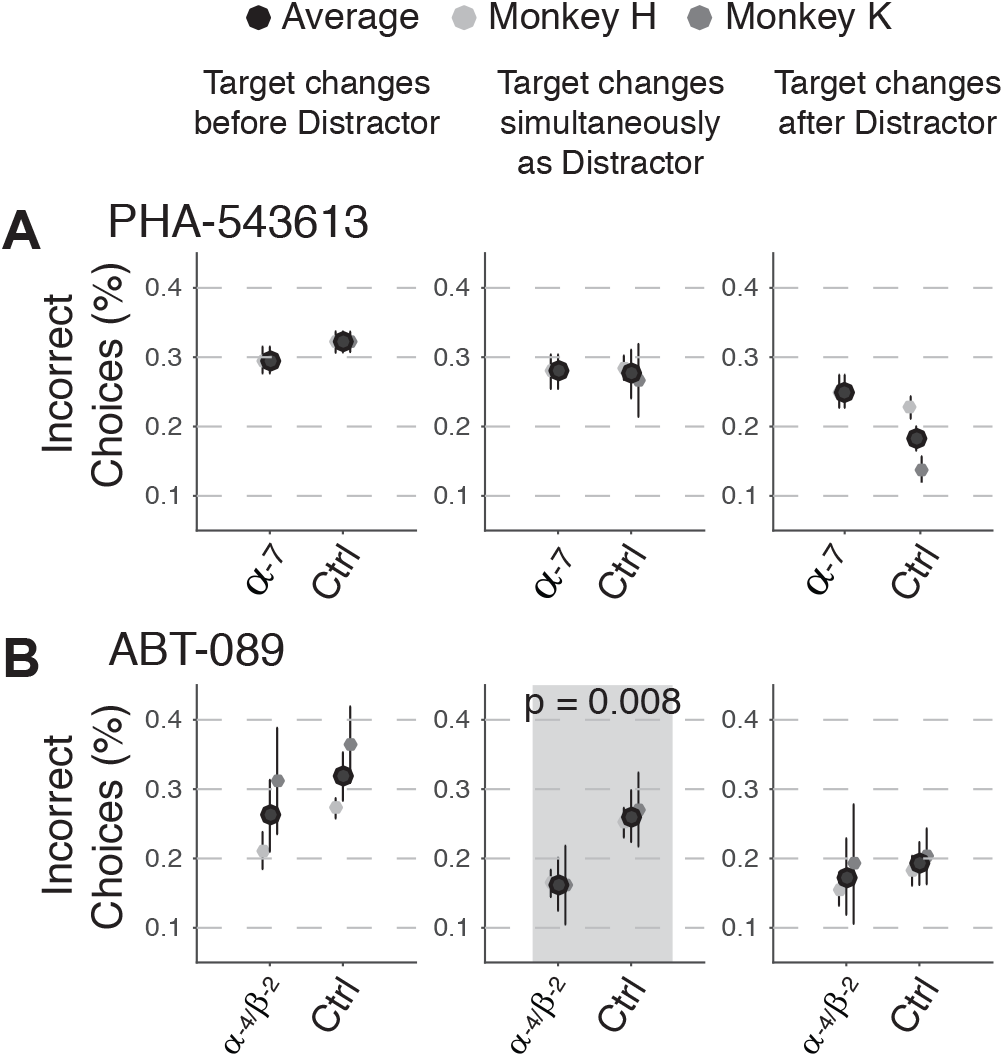
Sub-receptor specific effect on performance with different degrees of attentional filtering requirements. (**A**) Y-axis: Proportion of incorrect choices in trials when the Go/No-Go choice signal occurred in the target stimulus before the distractor (left column), at the same time as the distractor (middle column), and after the distractor (right column). The demand of attentional filtering of the distractor event is strongest when distractor and target change at the same time (middle column). The x-axis denotes the 0.125 mg/kg dose of the α7 agonist PHA-543613 and the control condition (saline injections). The black marker denotes the average across Monkey H (light grey) and Monkey K (dark grey). Error bars are standard errors of the mean. (**B**) Same format as (***A***) for the results obtained with the α4β2 agonist ABT-089. There was a significant improved performance across monkeys for the condition with enhanced attentional filtering requirements with the α4β2 agonist ABT-089. The x-axis denotes 0.02 mg/kg and 0.01 mg/kg doses for Monkey H and Monkey K respectively (Wilcoxon ranksum, p=0.008, grey shaded panel in A), but not in any other condition.

For the PHA-543613 treatments, the average performance accuracy in these conditions were indistinguishable from control performance at the 0.125 mg/kg and 0.25mg/kg conditions for monkey’s H and K (Wilcoxon ranksum tests for each dose and monkey, p>0.05) (**Fig. 3A**).

In contrast, ABT-089 treatment yielded lower errors than control when both stimuli dimmed simultaneously in both monkeys, but not in other trials types in which the target dimmed before or after the distractor dimmed (**Fig. 3B**). In Monkey H, ABT-089 treatment with 0.02 mg/kg led to reduced erroneous distractor choices in trials with simultaneous dimming than control treatment (Wilcoxon ranksum test, p=0.031). Drug versus control performance was similar for the other conditions with the dimming occurring before or after the distractor. There was no significant drug effect on performance with and without filtering demands with 0.01 mg/kg ABT-089 treatment. In Monkey K, ABT-089 treatment with 0.01 mg/kg there was a trend for reduced erroneous distractor choices in trials with simultaneous dimming (16% erroneous choices) than control treatment (27% erroneous choices) (Wilcoxon ranksum test, p=0.16) (**Fig. 3B**). There was no performance difference when the target dimmed before or after distractor (Wilcoxon ranksum test, n.s.). At the higher 0.02 mg/kg dose the performance was not significantly different than control for trials with filtering demands (Wilcoxon ranksum test, p=0.32).

The results showed the same direction of the ABT-089 drug effect to improve performance during trials with larger filtering demands, but not in other trial types in Monkey H at 0.02 mg/kg, and in Monkey K at 0.01 mg/kg. We therefore pooled the data from Monkey H ABT-089 (0.02 mg/kg) and Monkey K (0.01 mg/kg) to increase statistical power for inferential analysis. This pooling showed a highly significant difference between drug and control sessions only when both stimuli dimmed simultaneously (Wilcoxon ranksum, p=0.008) (black marker in **Fig. 3B**). No significant difference was observed with any other dimming conditions (Wilcoxon ranksum, p>0.1). Taken together, ABT-089, but not PHA-543613, was associated with improved attentional filtering.

## Comparison of Baseline Performance Levels

We tested for PHA-543613 and ABT-089 drug effects in experimental sessions that were separated by several weeks (see Methods). We therefore tested whether the overall performance levels remained similar between testing periods for either drug. To this end we extracted the control performance during the PHA-543613 testing sessions, and compared them to the ABT-089 testing sessions. We found that for both monkeys the asymptotic performance levels (the Rmax parameter of the hyperbolic ratio fits) were similar for PHA-543613 and ABT-089 testing sessions (randomization tests, all p>0.05). With regard to learning, we found that monkey H showed faster learning during the ABT-089 control sessions than in the PHA-543613 control sessions (randomization, p=0.0133). There was no difference in learning speed in control sessions for monkey K (randomization test, p>0.05). We performed the same control session comparison for the attentional filtering performance. We found for monkey K that control performance was statistically indistinguishable for the different conditions (target dimming before, after, or simultaneously with distractor) (randomization tests, all p>0.05). Monkey H showed similar performance in PHA-543613 and ABT-089 control testing sessions when the target dimmed before or simultaneously with the distractor, but was more accurate for in the ABT-089 control sessions when the target dimmed after the distractor (see Supplementary Fig. 3). In summary, these findings indicate that despite an overall consistent performance pattern in monkey K, monkey H showed selected improvements during the ABT-089 testing sessions.

## Effects of Selective nAChR on Error Types and Reaction Times

The effects of PHA-543613 on learning and the effects of ABT-089 on attentional filtering might be mediated by proneness for specific behavioral errors, or by trading accuracy with reaction times (speed-accuracy trade-offs). To address the first factor, we quantified three different types of errors. First, we considered perseveration errors, estimated as the proportion of successive unrewarded choices following a correct choice. Second, we counted the number of fixation breaks in the covert attention period that started with the onset of the stimulus color and motion, and ended with the dimming event prior to the choice. These errors might index lapses in attentional control to prevent looking at the peripheral stimuli. Third, we calculated the overall number of fixation breaks that occurred early in the trial, prior to the color and motion onset, which might indicate overall oculomotor tone or motivational factors. We found that the occurrences of neither of these three error types varied with PHA-543613, or ABT-089 at low, or high doses in Monkey H or Monkey K (all Wilcoxon ranksum tests comparing each drug condition to their control conditions, p>0.05).

To address possible drug effect on reaction time speed, we calculated the saccadic reaction times following the choice event. We found that none of the drugs resulted in statistical changes in reaction times at low or high doses in Monkey H or Monkey K (all pairwise Wilcoxon ranksum tests, p>0.2). Average reaction times across drugs and experimental conditions ranged from 188 ms to 197 in Monkey H, and from 232 to 247ms in Monkey K.

## Discussion

We found that two sub-receptor specific nicotinic agonists have dissociable effects on the learning speed and the attentional filtering performance of two healthy rhesus macaques. The selective α7 nAChR agonist PHA-543613 selectively enhanced the speed of learning feature values, but did not modulate how salient, distracting information was filtered from ongoing choice processes. In contrast, the selective α4β2 nAChR agonist ABT-089 did not affect learning speed but reduced the number of intrusions of distractor information in the condition that required the attentional filtering of a salient distractor event while maintaining the focus on a target stimulus. This double dissociation of learning versus filtering processes was evident for each monkey, and only for one of two drug doses tested. Moreover, these dose-dependent behavioral effects were evident in the absence of systematic changes in overall performance, reward intake, motivation to perform the task, perseveration tendencies, or reaction times. Overall, we believe that these findings provide strong evidence for sub-receptor specific nicotinic mechanisms supporting higher level attention and learning functions in the primate brains.

### α7 nAChR specific enhancement of cognitive flexibility to adjust to behavioral feedback

We found that at the higher dose, the α7 agonist gave rise to faster learning following feature-reward reversals in both monkeys (**Fig. 2**). The learning effect disappeared within 25 minutes and 50 minutes for Monkey H and K, respectively. To understand the temporal specificity of this effect we performed HPLC analysis of plasma concentration level temporal decay of PHA-543613 as an indirect estimate of neurally available drug concentration. We found that our monkeys started performance within minutes after the peak plasma concentration levels and that after 50 minutes of task performance the plasma concentration levels had dropped to below half maximum concentration level (**Figure S1**). We believe this rapid kinetic underlies the short-lived improvement on learning speed with the α7 agonist PHA-543613.

The observed faster reversal learning indexes enhanced flexibility to adjust to changing behavioral feedback and thus reflects a genuine improvement of attentional control and learning from error feedback (Kehagia AA et al. 2010; Balcarras M et al. 2016; Izquierdo A et al. 2017). Such enhanced flexibility can have multiple causes. Previous cholinergic modulation studies suggest that enhanced flexibility to adjust to behavioral challenges are mediated by an enhanced tonic mode of cholinergic action in prefrontal cortex (Sarter M *et al.* 2016). According to this model cholinergic transients are triggered by unexpected increases in performance demands. Enhanced ACh availability is then allowing α7 nAChRs action to support more robust neural goal representations. More robust goal representations translate into behavior that more effectively ‘follows-through’ to achieve the goal. A key observation of this model is that in rodent prefrontal (peri-and infralimbic) cortex ACh transients occur during a sustained attention task in correct trials that followed misses or correct rejections (Parikh V et al. 2007; Howe WM *et al.* 2013). These trials are similar to the post-reversal trials in our task in which the animal experiences unexpected omissions of reward after choices made to the no longer reward associated stimulus color (causing negative prediction errors), and in subsequent trials when the animals receive reward to the previously unrewarded stimulus color. Consequently, our results support the suggestion that stronger α7 receptor activation enhances the robustness of neural goal representations and thereby facilitating to “follow-through” in adjusting attentional set and behavior after a reward-reversal. This receptor specific interpretation is consistent with the finding in rodents that α7 nAChRs action prolongs the duration of elevated ACh to >10 sec following cholinergic transients in the prefrontal cortex (Parikh V *et al.* 2010). Such longer durations are likely beneficial in our task to successfully follow-through with adjusting the attentional set across multiple post-reversal trials, each lasting ~4-6 sec.

The neural effect underlying the proposed α7 action might well involve changes in synaptic connection weights among neurons encoding the newly rewarded stimulus as suggested by reversal learning models (Fusi S et al. 2007; Rombouts JO et al. 2015). Intriguingly, such short-term plasticity might be particularly supported by the α7 receptor which is associated specifically with enhanced intracellular calcium signaling, spike-timing dependent plasticity and long-term potentiation of synapses in medial temporal cortex and cortico-amygdala circuits (Gu Z and JL Yakel 2011; Gu Z et al. 2012; Jiang L et al. 2016). We believe that similar α7 mediated learning might be effective in fronto-striatal circuits that are critical for feature-based attention and reversal learning.

A possible complementary route for how α7 specific activity could support reversal learning in our task is by amplifying other neurotransmitter systems. Previous studies have shown that in prefrontal cortex, striatum and hippocampus α7-receptor activation triggers release of glutamate (Jones IW and S Wonnacott 2004; Dickinson JA et al. 2008; Bortz DM *et al.* 2013), GABA (Arnaiz-Cot JJ et al. 2008), norepinephrine (Kennett A et al. 2012), and dopamine (Pidoplichko VI et al. 1997; Livingstone PD and S Wonnacott 2009; Quarta D et al. 2009). Among these co-activated neurotransmitters, dopamine and norepinephrine have long been implicated to enhance cognitive flexibility in reversal learning and attentional set shifting tasks (van der Meulen JA et al. 2007; Kehagia AA *et al.* 2010; Wallace TL and D Bertrand 2013). Consequently, activating α-2A norepinephrine receptors could be one route α7 could improve learning. Strong support for this scenario comes from a previous study in which we found a similar faster reversal learning performance at optimal doses of the α-2A noradrenergic agonist Guanfacine (Hassani SA *et al.* 2017). In that study, an increased reversal learning speed was the most prominent behavioral effect during prolonged testing at effective doses.

Our study adds an important data point to the five existing studies testing selective nAChR’s agonists on set shifting or reversal performance in monkeys (**Table 1**, **Table’s S1**, **S2**) by illustrating that selective α7 nAChR activation specifically enhanced the learning speed without affecting other attentional markers of task performance. This effect was dose specific. A previous study reported detrimental set shifting effects of an α7 receptor agonist at a dose at which the agonist enhanced short term memory maximally (Gould RW *et al.* 2013). We believe that our dose specific finding suggests that activating α7 specific mechanisms can enhance learning flexibility, but that this effect occurs independently of strengthening recurrent short-term memory activity. Consistent with this interpretation it has been acknowledged that strong short-term memory that might be needed for bridging long delays in delayed match-to-sample tasks conflicts with the demands to flexibly shift between attended stimulus features needed to succeed with reversal learning (Thiele A and MA Bellgrove 2018).

To summarize, our finding of dose specific enhanced reversal learning is consistent with α7 specific modulation of the duration of elevated cholinergic tone following cholinergic transients putatively triggered by reward reversals. The working mechanism might involve α7 mediated short-term strengthening of synaptic connections in prefrontal cortex or striatum but may also include the amplification of other neurotransmitter actions, particularly the norepinephrinergic receptors.

### α4β2 specific filtering of distraction and stabilizing of goal representations

Our task varied the attentional filtering demands and found that the α4β2 agonist, but not the α7 agonist, facilitated attentional filtering of a distractor event occurring at the same time as the choice event of the target stimulus (**Fig. 3**). ABT-089, but not PHA-543613, reduced the choice errors by half compared to the control condition, strongly suggesting that enhanced distractor filtering is specifically mediated by α4β2 mechanisms. Prior NHP studies demonstrated distractor filtering benefits with α4β2 agonists using delayed match-to-sample tasks (see **Table 2** and **Table S2**). However, these NHP studies did not include α7 nAChR modulation to allow a direct comparison between sub-receptor specific effects with the same subjects and experiment. Our receptor-specific dissociation is consistent with suggestions of α4β2 sub-receptor specific distractor modulation in rodents using a sustained attention task (SAT) (Howe WM *et al.* 2010). The SAT requires rodents to memorize a cue location and either respond or withhold responding to that location. Distracting light flashes between cue and response intervals are more tolerated during α4β2 agonist action than α7 agonist action (Howe WM *et al.* 2010; St Peters M *et al.* 2011). Our study extends these insights in receptor-specific facilitation of distractor filtering to the NHP domain and a feature-based attention paradigm.

**Table 2.** Overview of behavioral results from systemic applied nAChR on the delayed-match-to-sample task with and without distracting events during the delay (DMTS versus dDMTS), and on the sustained attention task with and without delay (SAT, dSAT). “+”: positive effect on performance; “-”: negative effect on performance; “N”: no effect. The details about the included studies are outlined in **Table S1** and **Table S2**.

|  |  | DMTS | dDMTS | SAT | dSAT |
| --- | --- | --- | --- | --- | --- |
| $\alpha 4\beta 2^*$ | NHP: | +*1N +++++ | +++ | | |
|  | Rat: |  |  | + - | + |
| $\alpha 7^*$ | NHP: | +++*2+*3++ | +*4 | | |
|  | Rat: |  |  | + |  |

\* $\alpha 4\beta 2$ agonists included ABT-418 ( $\alpha 4\beta 2$ agonist), ABT-594 ( $\alpha 4\beta 2$ agonist), S-38232 ( $\alpha 4\beta 2$ agonist), SIB-1508Y ( $\alpha 4\beta 4$ agonist); $\alpha 7$ agonists included RG-3487 ( $\alpha 7$ agonist), A-582941 ( $\alpha 7$ agonist), PHA-543613 ( $\alpha 7$ agonist)
\*1: ABT-089 ( $\alpha 4\beta 2$ agonist, young and aged NHP, slow and optimal learner rats),
\*2: GTS-21 ( $\alpha 7$ agonist, healthy and ketamine impaired monkeys)
\*3: PNU-282987 ( $\alpha 7$ agonist, cocaine-experienced & naïve),
\*4: Varenicline (agonist for $\alpha 4\beta 2$ & $\alpha 7$ , ketamine impaired monkeys only)

In fact, we are not aware of other NHP paradigms that succeeded to isolate a distractor-filtering specific drug effect in the absence of confounding short-term memory manipulations. We believe this is a significant advance of the field. Despite the similarity of the functional benefits of α4β2 action in light of attentional filtering demands in the DMTS (in NHP and rodents), the SAT (in rodents), and the feature-based attentional learning task we deployed, there are important differences of the nature of the distracting events that are affected in these different tasks. Firstly, prior NHP studies using ABT-089 indicate that flashing distracting random lights in the first 3s of the delay of a delayed match to sample task are not impairing behavior when the matching stimulus follows ~10 sec. after the cue (Prendergast MA, WJ Jackson*, et al.* 1998). This finding indicates that with good dosing of α4β2 agonists the cue (the DMTS sample stimulus) is more robustly represented in short-term memory, as has been directly documented in prefrontal cortex neurons (Sun Y et al. 2017). The described distractor filtering during short term memory delays in these tasks is similar to the attentional filtering condition in our task in which the distractor stimulus undergoes a luminance change just prior the target choice event, i.e. when the target representation had to be selectively maintained while a single salient event distracted ongoing attention. However, in contrast to the prior DMTS NHP studies (Prendergast MA, WJ Jackson*, et al.* 1998; Buccafusco JJ *et al.* 2007), in our ‘target after distractor event’ condition neither the α4β2, nor α7 agonist improved performance of our animals. We believe that we did not find a distractor effect in this condition because the subjects did not need to rely on a purely internally maintained short-term memory of the target feature (the rewarded color) but could refocus attention to the target stimulus even when the distractor captured attention. Thus, α4β2 agonists could not reveal its role in enhancing the internal goal representation in this condition, as long as there was sufficient inhibitory behavioral control to not respond to the distracting event when it occurred before the choice event in the target stimulus.

We did find, however, that good dosing of α4β2 agonists enhanced filtering distracting events when they co-occurred with the target event. Such simultaneous perceptual events are the hallmarks of perceptual conflict studies of cognitive control (Alexander WH and JW Brown 2011) and to our knowledge have not been tested in NHPs with selective nAChR agonists. We believe the most parsimonious explanation for this effect is the selective stabilization of goal – relevant feature representations as recently suggested in the framework of neurotransmitter control of neural attractor dynamics (Thiele A and MA Bellgrove 2018). In this framework, increased distractor filtering follows from stronger recurrent dynamic activity of neural circuits representing the attended target feature. Consistent with this scenario, active α4β2, but not α7, nAChRs have been shown to gain modulate the amplitude of phasic cholinergic increase in prefrontal cortex when encountering unexpected outcomes (Parikh V *et al.* 2007; Parikh V *et al.* 2010).

Taken together, our findings provide strong converging evidence for the assertion that α4β2 acts as gain modulator for the strength of prefrontal cholinergic transient activity, while α7 receptor activation mediates longer durations of cholinergic tone in prefrontal cortex (Parikh V *et al.* 2010). We believe that future studies could test this model by measuring the influence of selective nAChR‘s on neural activity dynamics in the prefrontal cortex and associated network nodes in the striatum.

### Implications of a functional double dissociation of α7 and α4β2 receptor action on learning and attention

During natural behavioral without neurochemical interventions, both α7 and α4β2 receptor subtypes are activated when acetylcholine becomes available through stimulation of the medial forebrain bundle (Ballinger EC *et al.* 2016). By applying systemic nAChR agonists we have biased the natural balance of α7 and α4β2 action towards either sub-receptor action. By revealing a functional double dissociation of biasing the balance to either α7 or α4β2 we believe our findings reveal that these sub-receptors tap into distinct operations in the neural circuit that have been underestimated in previous studies. A recent comprehensive survey of cholinergic modulation of attention concluded that α7 and α4β2 actions might tap into similar functionality to stabilize goal representations in the prefrontal cortex, but that α7 receptor action is less potent than α4β2 action in the PFC than in memory related circuits (Thiele A and MA Bellgrove 2018). While our finding is consistent with this view, it raises the possibility that both sub-receptor systems play fundamentally different roles in neural circuits that are camouflaged by their co-modulation in most natural conditions and by their common action to trigger NMDA dependent increases in excitability (Yang Y *et al.* 2013; Sun Y *et al.* 2017).

In addition to the alleged distinct roles within the same (prefrontal cortex) circuit, the observed double dissociation would also be consistent with a scenario in which α7 receptor action and α4β2 affect different circuits, each contributing uniquely to learning and to attentional filtering (Chudasama Y and TW Robbins 2006). We believe that this suggestion is unlikely explaining the observed functional dissociation, given that both sub-receptors are widely available across all major cortical and subcortical systems relevant for attentional control and reversal learning (**Figure S4**) (Gotti C *et al.* 2006). However, it might soon be possible to discern more subtle sub-receptor density differences with recent advances in whole brain PET imaging of α7 and α4β2 receptor availability (Teodoro R et al. 2015; Hillmer AT et al. 2017).

Such differences in sub-receptor specific expression density of α7 and α4β2 nAChRs have been a hallmark of studies in clinical populations. A most prominent example is schizophrenia, where patients have been repeatedly found to show reduced α7 receptor expression and availability in anterior cingulate cortex, frontal cortex and hippocampus (Martin-Ruiz CM et al. 2003; Wong DF et al. 2018). Consistent with these findings, α7 is implicated in the pathophysiology and the cognitive inflexibility symptoms of schizophrenia, as well as of Alzheimer’s diseases (Thomsen MS et al. 2010; Toyohara J and K Hashimoto 2010). Our findings are remarkably consistent with these findings of α7 associated behavioral flexibility, and suggest that the NHP animal model will be essential to understand the underlying mechanisms of altered nAChR functioning in schizophrenia and Alzheimer’s dementia.

## Conclusion

Our results document a double dissociation of cholinergic sub-receptor specific contributions to higher-order learning and attention functions in the non-human primate. We believe that these findings will reinforce ongoing efforts in developing drugs for alleviating the symptoms in schizophrenia and Alzheimer’s, as well as in ADHD and various other neuropsychiatric conditions in which behavioral flexibility and attentional focusing is compromised together with altered α7 and α4β2 signaling (Millan MJ et al. 2012; Bertrand D et al. 2015).

## Acknowledgments

This work was supported by a grant from the Canadian Institutes of Health Research, CIHR Grant MOP_102482 (T.W.). The funders had no role in study design, data collection and analysis, the decision to publish, or the preparation of this manuscript. The authors would like to thank Hongying Wang for technical support.

---

**Figure S1.**
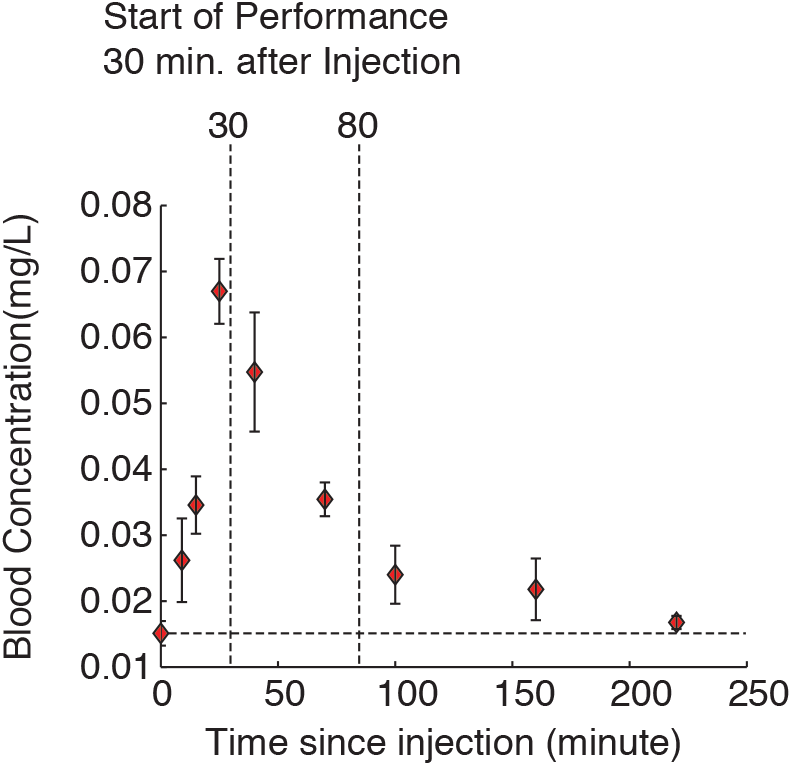
Measured PHA-543613 plasma exposure (mean ± standard deviation) in Monkey S. HPLC analysis of blood serum obtained over a period of 220 minutes after IM injection of 0.250 mg/kg dose.

**Figure S2.**
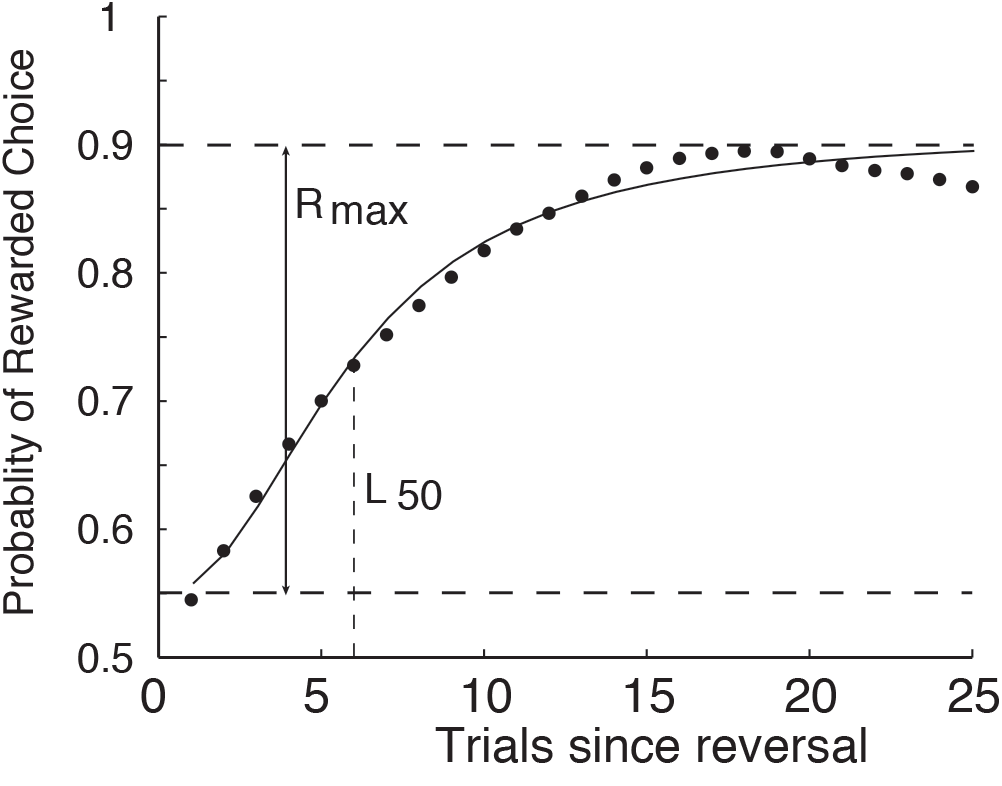
Parameters of hyperbolic ratio function fit to an example learning curve. The L_50_ parameter indexes the trial at which the animal reached half maximal performance (in this example, trial 6). The R_max_ parameter is the difference between baseline and maximal performance indicated by lower and upper dash lines respectively.

**Figure S3.**
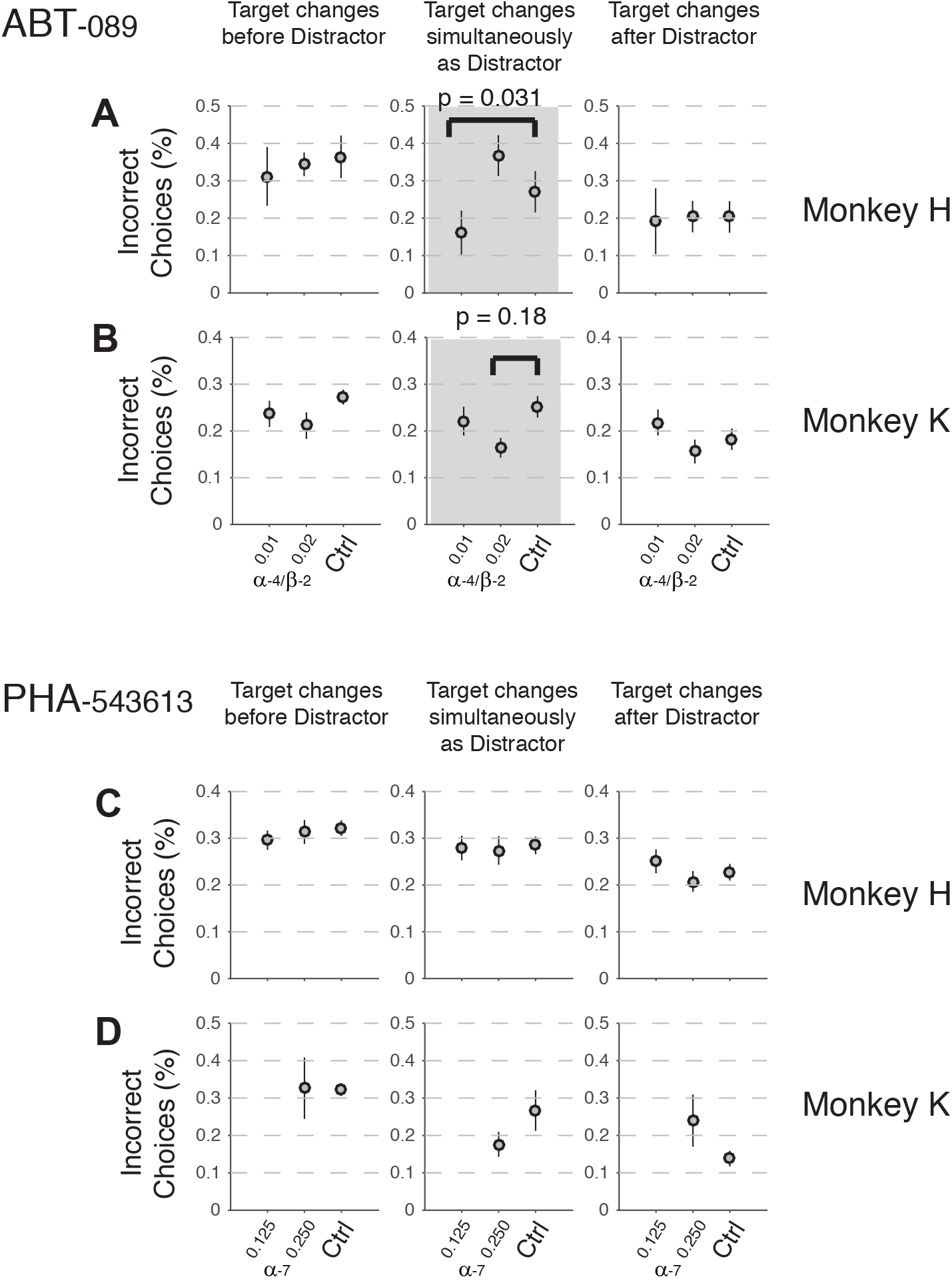
Average Performance with different degrees of attentional filtering requirements for each monkey, drug, and dose. (**A,B**) For ABT-089, proportion of incorrect choices in trials when the Go/No-Go choice signal occurred in the target stimulus before the distractor (left column), at the same time as the distractor (middle column), and after the distractor (right column). The demand of attentional filtering of the distractor event is strongest when distractor and target change at the same time (middle column). The x-axis denotes the different drug and control conditions. Results are shown separately for Monkey H (**A**) and Monkey K (**B**), respectively. Error are standard errors of the mean. (**C,D**) Same format as (A,B) for the α7 agonist PHA-543613. (Wilcoxon ranksum, p=0.031).

**Figure S4.**
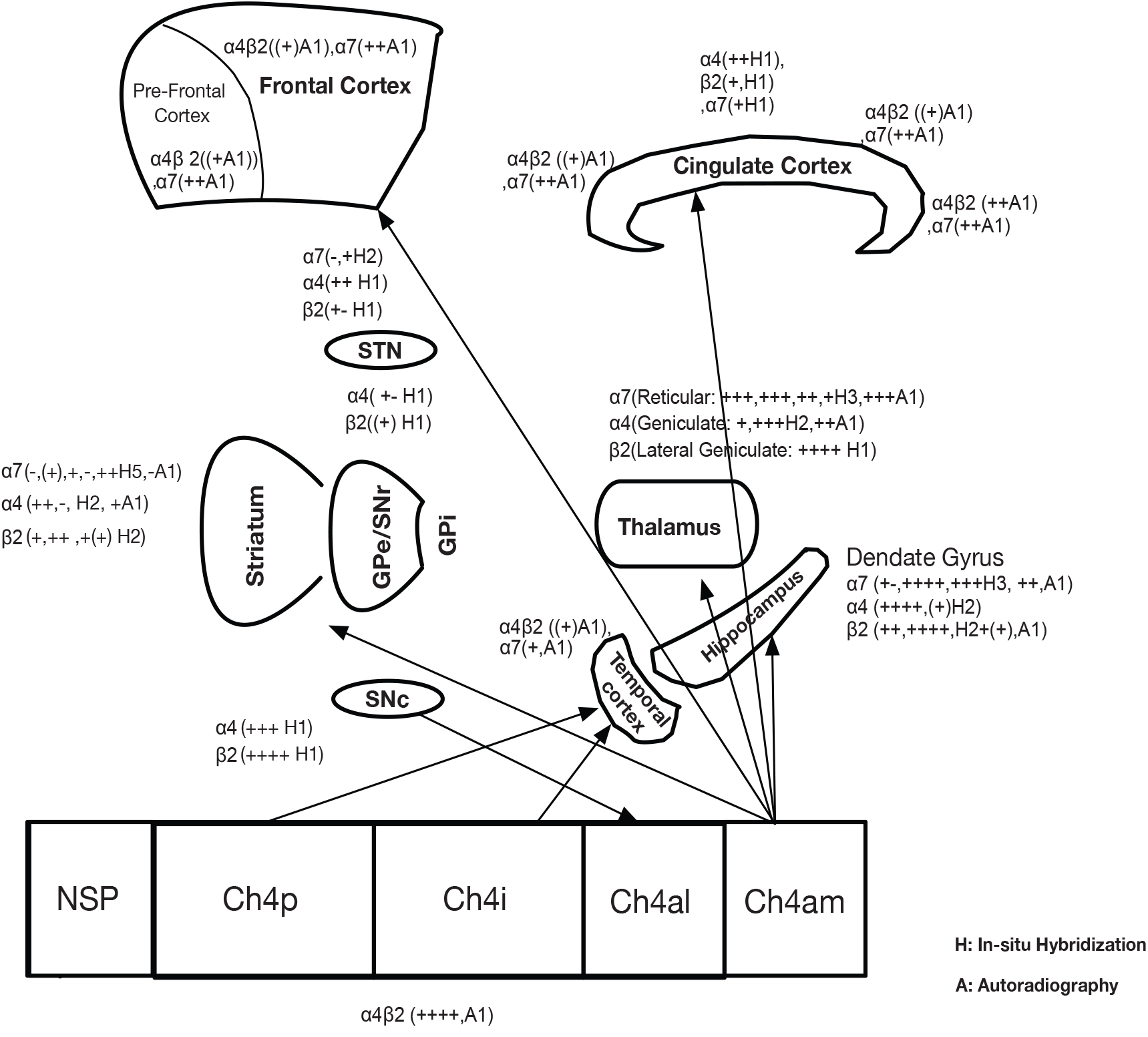
Suggestive localization of nACh receptors in cortical and sub-cortical areas based on results from human and non-human primate studies. These results were obtained via in-situ hybridization and immunocytochemistry techniques (Cimino et al., 1992; Kulak et al., 2006; Quik et al., 2000; Quik et al., 2005; Spurden et al., 1997; Han et al., 2000; Han et al., 2003).

## Footnotes

1 mean ± standard error of mean

